# Improved inter-protein contact prediction using dimensional hybrid residual networks and protein language models

**DOI:** 10.1101/2022.08.04.502748

**Authors:** Yunda Si, Chengfei Yan

## Abstract

The knowledge of contacting residue pairs between interacting proteins is very useful for structural characterization of protein-protein interactions (PPIs). However, accurately identifying the tens of contacting ones from hundreds of thousands of inter-protein residue pairs is extremely challenging, and performances of the state-of-the-art inter-protein contact prediction methods are still quite limited. In this study, we developed a deep learning method for inter-protein contact prediction, referred to as DRN-1D2D_Inter. Specifically, we employed pretrained protein language models to generate structural information enriched input features to residual networks formed by dimensional hybrid residual blocks to perform inter-protein contact prediction. Extensively benchmarked DRN-1D2D_Inter on multiple datasets including both heteromeric PPIs and homomeric PPIs, we show DRN-1D2D_Inter consistently and significantly outperformed two state-of-the-art inter-protein contact prediction methods including GLINTER and DeepHomo, although both the latter two methods leveraged native structures of interacting proteins in the prediction, and DRN-1D2D_Inter made the prediction purely from sequences.

## Introduction

Structural characterization of protein-protein interactions (PPIs) is of great importance for mechanistic investigation of cellular processes and therapeutic design^1–3^. However, currently we still lack structural information for most of known PPIs, as determining complex structures of PPIs using techniques like X-ray crystallography, cryogenic electron microscopy (cryo-EM) is time consuming and expensive^4–6^. The knowledge of contacting residue pairs between interacting proteins is very useful for structural characterization of PPIs^7–10^. Therefore, the prediction of residue-residue contacts between interacting proteins is of great significance. However, accurate inter-protein contacts prediction is extremely challenging, as a protein-protein interaction formed by proteins with average sizes (e.g. ∼300 sequence length) generally have hundreds of thousands of inter-protein residue pairs (e.g. ∼300*300), but only tens of them form contacts (e.g. heavy atom distance < 8.0 Å) in 3D space. Previous studies for inter-protein contact prediction are often based on the fact that contacting residue pairs often vary cooperatively in the process of evolution, thus in which coevolutionary analysis methods were used to predict inter-protein contacting residue pairs^7,8,11–14^. However, coevolutionary analysis-based inter-protein contact prediction methods do have certain limitations. For example, for homomeric PPIs (i.e. the two interacting proteins are the same protein), the inter-protein coevolutionary signals are mixed with the intra-protein coevolutionary signals, thus it is difficult to tell whether the sequence covariation is caused by inter-protein contact or intra-protein contact^13^. For heteromeric PPIs (i.e. a protein interacts with a different protein), it is often difficult to accurately identify interologs due to the existence of paralogs^7,8^. However, building the multiple sequence alignment (MSA) of interologs is required for coevolutionary analysis. Besides, since effective coevolutionary analysis generally requires a large number of sequences, for many PPIs, even though their interologs can be correctly identified, the number of existing interologs may not be enough to guarantee the effectiveness of coevolutionary analysis. Due to the limitations above, the coevolutionary analysis-based inter-protein contact prediction methods only work well for a very limited number of PPI cases.

Apart from co-evolutionary analysis methods, deep learning techniques have also been applied to predict inter-protein contacts for homomeric and/or heteromeric PPIs, inspired by their great successes in intra-protein contact prediction^15–18^. These methods generally employed inter-protein coevolutionary information obtained from coevolutionary analysis as one of the major input features to deep residual networks, the deep learning architecture which dominants the intra-protein contact prediction. For example, Zeng et al. developed ComplexContact, to the best of our knowledge, the first deep learning method for inter-protein contact prediction for heteromeric PPIs^15^. ComplexContact directly leveraged deep learning models trained for intra-protein contact prediction to perform inter-protein contact prediction, in which the input features including evolutionary and co-evolutionary information were drawn from two MSAs of interologs prepared by a genome-based approach and a phylogeny-based approach respectively. ComplexContact significantly outperforms co-evolutionary methods for inter-protein contact prediction, however, the overall performance is still quite limited, especially for PPIs in eukaryotes, as accurately identifying interologs in eukaryotes is much more challenging. Comparing with heteromeric PPIs, the co-evolutionary information of homomeric PPIs can be obtained much more easily, as each homologous protein interacts with itself to form the interolog. However, since both intra- and inter-protein contacts can cause sequence covariations in evolution, it is often difficult to determine whether the predicted contacts to be inter-protein or intra-protein. Yan et al. developed DeepHomo, a deep learning model specifically for inter-protein contact prediction for homomeric PPIs, in which they leveraged docking maps and distance maps drawn from protein monomeric structures as input features to a residual network to assist the identification of inter-protein contacts^18^. DeepHomo also significantly outperforms co-evolutionary analysis-based methods. However, DeepHomo requires high quality monomeric models as the input, which are not always available. Very recently, coming from the same group as ComplexContact, Xie et al. developed GLINTER^17^, a deep learning method which can perform inter-protein contact prediction for both heteromeric and homomeric PPIs. GLINTER first applied graph convolutional networks to learn rotational invariant representations for the monomeric structure of each interacting protein, which were then concatenated with the inter-protein attention maps generated by MSA transformer^19^ to form the input features to a residual network to predict inter-protein contacts. GLINTER achieved the-state-of-the-art performance in inter-protein contact prediction for both heteromeric and homomeric PPIs. However, GLINTER also requires monomeric models as the input, and according to its test result, the performance of GLINTER is quite sensitive to the quality of the inputted models.

In this study, we developed a new deep learning model for inter-protein contact prediction. Different from previous methods, we did not directly extract structural features from the monomeric structures but employed protein language models to generate structural information enriched features including 1D protein embeddings and 2D inter-protein attentions for each target PPI. The structural information enriched features obtained from protein language models were further combined with traditional statistical features including position-specific scoring matrices (PSSM) and inter-protein coevolution to form the input features of our prediction model, which were then transformed by the residual network formed by dimensional hybrid residual blocks to predict inter-protein contacts. It was shown in our previous study that the application of the dimensional hybrid residual blocks can increase the effective receptive field of the residual network, and thus improve the model performance^20^. The developed model referred to as DRN-1D2D_Inter was extensively benchmarked on PPI dataset including both homomeric PPIs and heteromeric PPIs. The result shows that DRN-1D2D_Inter significantly and consistently outperforms the two state-of-the-art methods DeepHomo and GLINTER for inter-protein contact prediction for both homomeric PPIs and heteromeric PPIs. It is worth noting that both the two reference models leveraged native structures of the interacting proteins in the inter-protein contact prediction, and DRN-1D2D_Inter made the prediction purely from sequences.

## Results

### 1. Overview of DRN-1D2D_Inter

#### 1.1 The model of DRN-1D2D_Inter

The model of DRN-1D2D_Inter is illustrated Figure 1. As it is shown in the figure, the input features of DRN-1D2D_Inter can be grouped into four categories: the 1D sequential features drawn from sequences of the interacting proteins, the 1D sequential features drawn from MSAs of homologous sequences of the interacting proteins, 2D inter-protein pairwise features drawn from the paired sequence (i.e. the concatenation of the two sequences) and 2D inter-protein pairwise features drawn from the paired MSA (i.e. the MSA of interologs). Specifically, given a PPI formed by two interacting proteins with sequence lengths as *L*_*A*_ and *L*_*B*_ respectively, the 1D sequential features drawn from sequences of the interacting proteins include embeddings of the two sequences from ESM-1b^21^, for which (*L*_*A*_,1280) and (*L*_*B*_,1280) are the dimensions of the embeddings respectively. The 1D sequential features drawn from MSAs for the two interacting proteins include the position-specific scoring matrices (PSSM) (with (*L*_*A*_,20) and (*L*_*B*_,20) as the dimensions respectively) and MSA embeddings outputted from ESM-MSA-1b^19^ (with (*L*_*A*_,768) and (*L*_*B*_,768) as the dimensions respectively). The 2D inter-protein pairwise features drawn from the paired sequence include the 660 inter-protein attention maps generated by ESM-1b ((*L*_*A*_, *L*_*B*_, 660)). The 2D inter-protein pairwise features drawn from the paired MSA include the 144 inter-protein attention maps generated by ESM-MSA-1b ((*L*_*A*_, *L*_*B*_, 144)), the inter-protein evolutionary coupling matrix calculated by CCMpred^22^, inter-protein mutual information matrix, average product correction (APC)-corrected mutual information matrix and contact potential matrix calculated by alnstats provided in MetaPSICOV^23^ ((*L*_*A*_, *L*_*B*_, 4)) (i.e. the inter-protein coevolution features). Before inputting to the network of DRN-1D2D_Inter, all the 1D sequential features of each interacting protein are first stitched together to form the 1D representations of the interacting proteins with dimensions (*L*_*A*_,2068) and (*L*_*B*_,2068) respectively, which are then converted into 2D pairwise feature maps with dimension ((*L*_*A*_, *L*_*B*_, 4136)) through outer concatenation. Finally, the 2D converted feature maps are combined with all other inter-protein 2D features to form the 2D input feature maps of DRN-1D2D_Inter ((*L*_*A*_, *L*_*B*_, 4944)).

**Figure 1.**
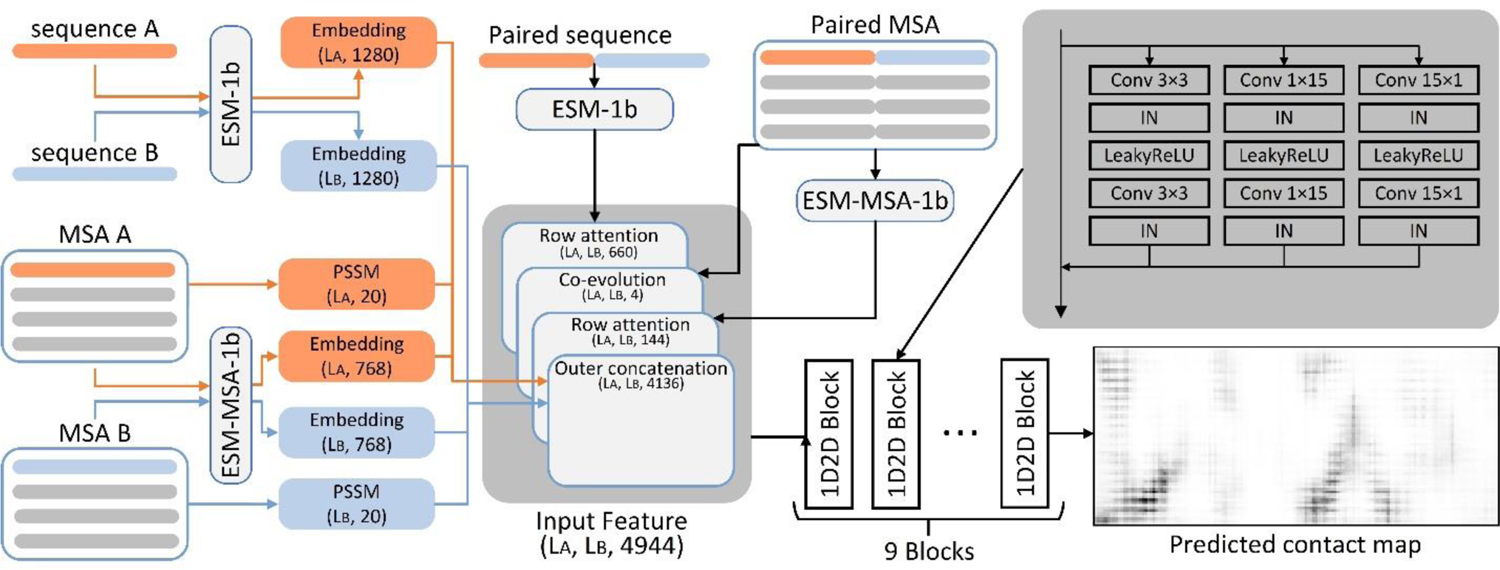
The network architecture of DRN-1D2D_Inter.

The input features are then transformed by the dimensional hybrid residual network formed by dimensional hybrid residual blocks (1D2D blocks). A detailed description of the 1D2D block can be found in (Si and Yan, 2021)^20^. In this study, we increased the length of the 1D convolution kernels from 9 to 15, for we found using the longer 1D kernels slightly improved the model performance. At the beginning of the network, a convolution layer with kernel size 1*1 is used to reduce the number of channels of the input features from 4944 to 96. After nine 1D2D blocks, another convolution layer with kernel size of 1*1 is used to transform the number of channels from 96 to 1, and then a sigmoid layer is applied to produce the predicted inter-protein contact map.

#### 1.2 The training protocol

For the development of DRN-1D2D_Inter, a non-redundant PPI dataset including 4828 homomeric PPIs and 3134 heteromeric PPIs was first prepared. Then we randomly selected 400 homomeric PPIs (denoted as ‘HomoPDB’) and 200 heteromeric PPIs (denoted as ‘HeteroPDB’) to form the test sets for the two types of PPIs respectively, and the remaining 4428 homomeric PPIs and 2934 heteromeric PPIs were used to tune the model.

To tune the model of DRN-1D2D_Inter, the 7362 PPIs (4428 homomeric PPIs and 2934 heteromeric PPIs) were equally split into seven subsets, each time we selected six subsets for the model training, and the left subset was used for the model validation. Seven independent models were trained separately, and the final prediction was the average of the predictions from the seven models. We used the singularity enhanced loss function proposed in (Si and Yan, 2021)^20^ to calculate training loss and optimized the loss with AdamW optimizer with its default settings. During training, if the loss on the validation set did not drop within two epochs, the learning rate would decay to 0.1 of its original (the initial learning rate was 0.001). The training would stop when the learning rate decayed twice and the model with the highest top-50 mean precision on the validation dataset was saved as the prediction model. The model was built with pytorch(v1.9.0) and trained on one NVIDIA TESLA A100 GPU with batch size equal to 1. Because of the memory limitation of GPU, the sequence length of each interacting protein was limited to 400. When a sequence was longer than 400, a fragment with sequence length equaling to 400 was randomly selected in the model training.

#### 1.3 The model evaluation

DRN-1D2D_INTER was first evaluated on HomoPDB and HeteroPDB, the two self-built datasets separated from our non-redundant PPI dataset. Apart from HomoPDB and HeteroPDB, two additional datasets including the homomeric PPIs from the test set of DeepHomo and the heteromeric PPIs from the Protein-protein Docking Benchmark 5.5^24^ were also used to evaluate the model performance. To make an unbiased evaluation, we removed PPIs if they have similar sequences to any PPI in our training dataset using 40% as the sequence similarity threshold, which yielded a dataset of 130 homomeric PPIs (denoted as DHTest) and a dataset of 59 heteromeric PPIs (denoted as DB5.5) respectively. In this study, two residues are defined to be in contact if the heavy atom distance of the two residues is smaller than 8 Å.

The two state-of-the-art models GLINTER and DeepHomo were also evaluated on the same datasets for comparison. Since DeepHomo is only developed for homomeric PPIs, it was only evaluated on HomoPDB and DHTest. Both GLINTER and DeepHomo require structures of the interacting proteins as the input to predict the inter-protein contacts. In this study, the bound structures extracted from the complex structure of each PPI were used as the input structures of the two models. It should be noted that in the real practice bound structures are generally not exist, and only unbound structures or even predicted structures can be used to assist the prediction. Therefore, the performances of GLINTER and DeepHomo can be overestimated with the application of bound structures in this study. Besides, it is worth mentioning that all the four test sets are not redundant with our training set, but there might be some redundancies between the four test sets and the training sets of GLINTER and DeepHomo, which may also cause the overestimation of the performances of GLINTER and DeepHomo.

In principle, the three tools (CCMpred, ESM-1b and ESM-MSA-1b) used to generate the input features of DRN-1D2D_Inter can also be directly employed to predict the inter-protein contacts. To show the effectiveness of our model, the inter-protein contact prediction performances of the three tools on the four test sets were also included for reference.

### 2. The performance of DRN-1D2D_Inter on HomoPDB and HeteroPDB

DRN-1D2D_Inter was first evaluated on the two self-built test sets: HomoPDB and HeteroPDB. Besides, the performances of GLINTER, CCMpred, ESM-1b and ESM-MSA-1b on the two test sets and the performance of DeepHomo on HomoPDB were also evaluated for comparison. We did not evaluate the performance of DeepHomo on HeteroPDB, as it was only designed to predict inter-protein contacts for homomeric PPIs. In Table 1, we show the mean precisions of the top (5, 10, 50, L/10, L/5) predicted contacts by the six methods on the two test sets, where L is the sequence length of the shorter protein in the PPI (Note: GLINTER failed to produce prediction results for 81 targets in HomoPDB and 15 targets in HeteroPDB, thus we removed these targets in the evaluation of the performance of GLINTER. The performances of other methods on HomoPDB and HeteroPDB after the removal of these targets are shown in Table S2). As it can be seen from the table that DRN-1D2D_Inter consistently outperformed all other methods on both test sets. Meanwhile, we can also see that CCMpred, ESM-1b and ESM-MSA-1b yielded much lower performances than DRN-1D2D, DeepHomo and GLINTER. It is reasonable since the latter three methods were tuned specifically for inter-protein contact prediction. Therefore, we focus on DRN-1D2D_Inter, DeepHomo and GLINTER for further analysis.

**Table 1.**
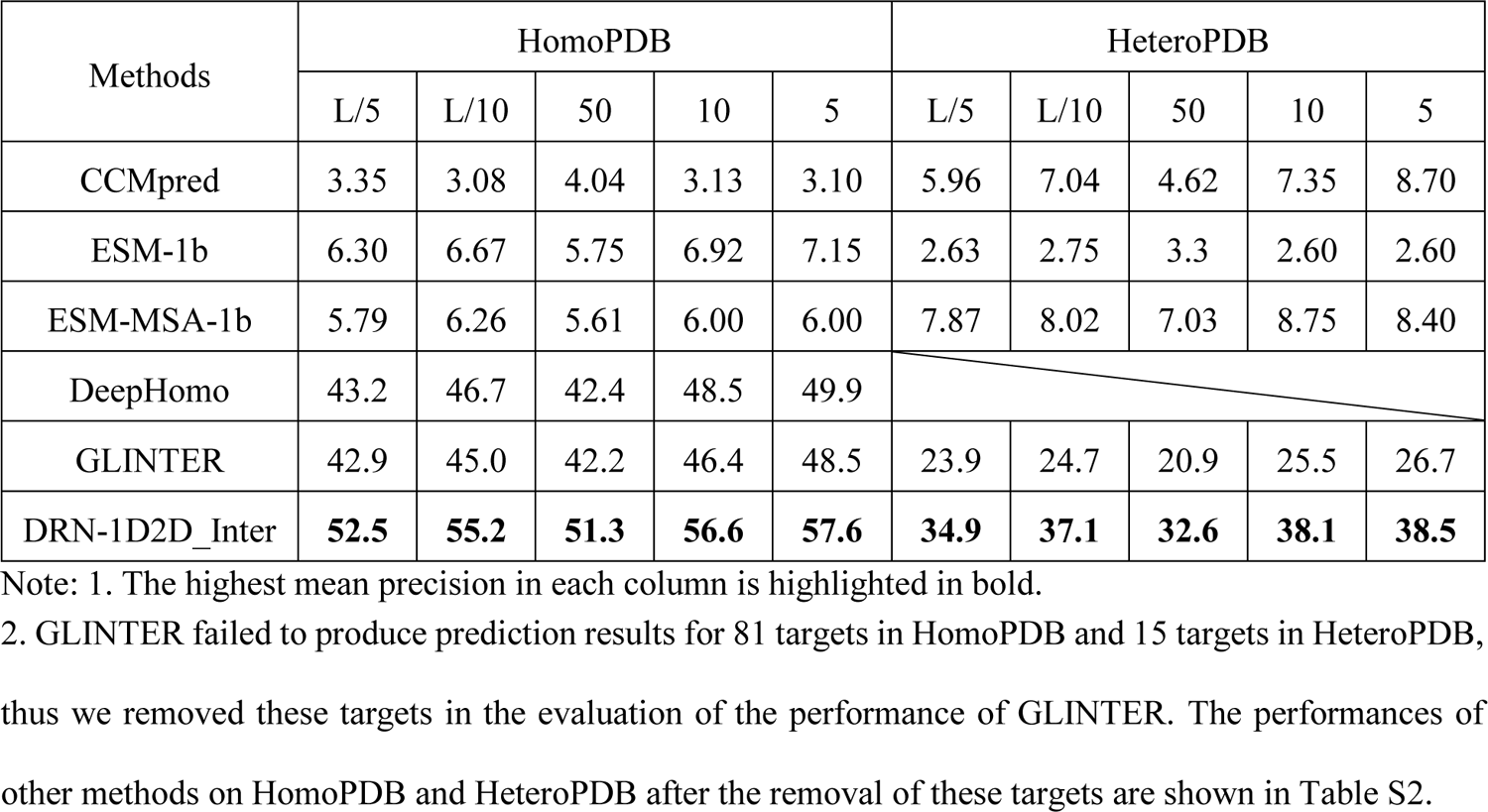
The performances of CCMpred, ESM-1b, ESM-MSA-1b, DeepHomo, GLINTER and DRN-1D2D_Inter on the HomoPDB and HeteroPDB test sets.

In Figure 2, we show statistical analysis of precisions of the top 50 predicted contacts by the three methods. Specifically, the violin plots of precisions of the top 50 predicted contacts by each method on the two datasets are shown in Figure 2a-b. From Figure 2a-b we can clearly see that DRN-1D2D_Inter significantly outperformed DeepHomo (p-value: 9.71e-4) and GLINTER (HomoPDB p-value: 9.87e-15, HeteroPDB p-value: 1.51e-8). We also show the complementary cumulative distribution function (CCDF) of precisions of the top 50 predicted contacts by each method on the two datasets are shown in Figure 2c-d. From Figure 2c-d we can see that on HomoPDB dataset, 55% of the targets predicted by DRN-1D2D_Inter have a precision higher than 50%, meanwhile only 41% and 43% of the targets predicted by DeepHomo and GLINTER have an accuracy above 50%; on HeteroPDB dataset, 28% of the targets predicted by DRN-1D2D_Inter have an accuracy above 50%, and only 18% of the targets predicted by GLINTER have an average accuracy above 50%.

**Figure 2.**
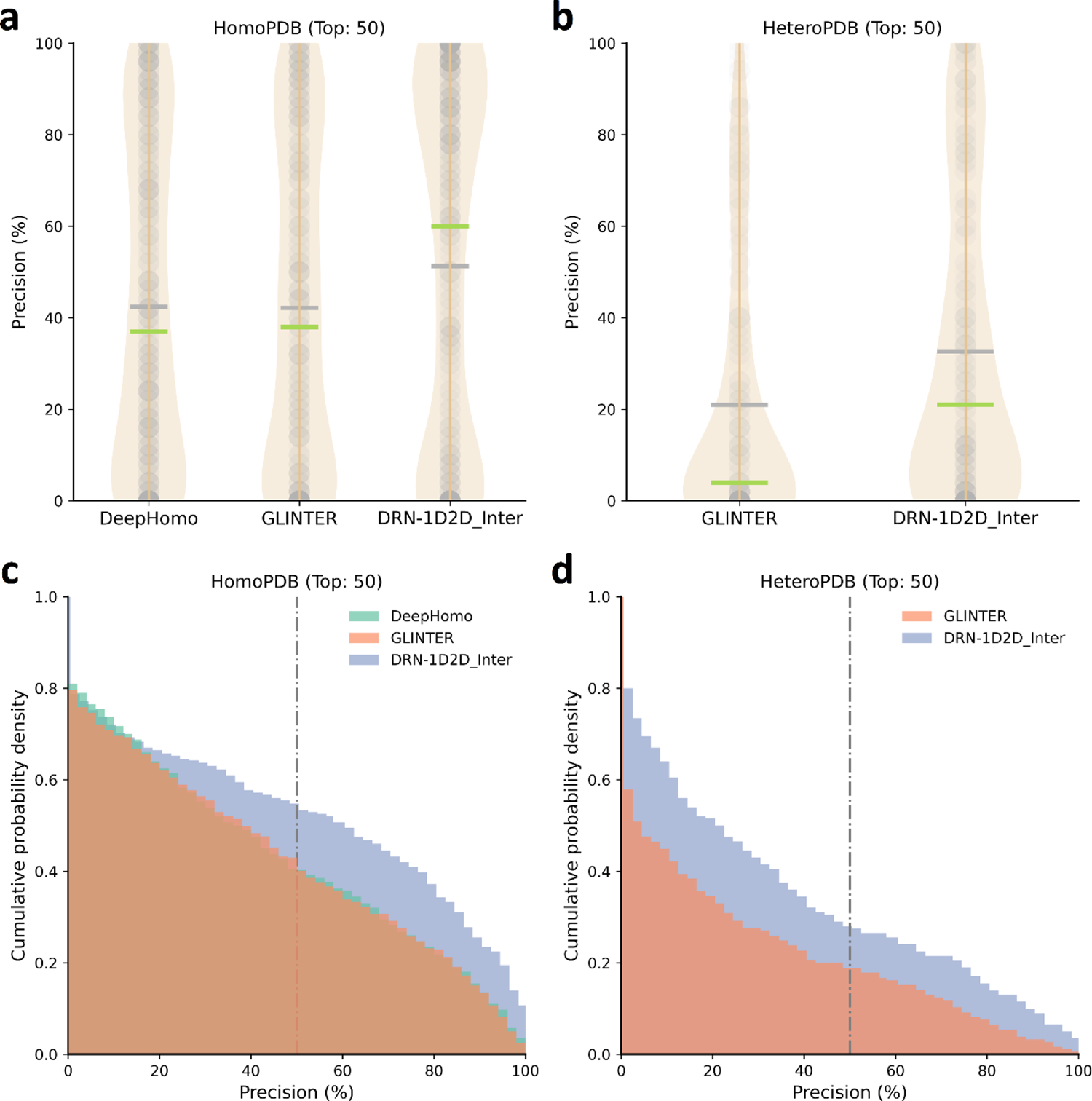
The statistical analysis of precisions of the top 50 predicted contacts by different methods on the HomoPDB and HeteroPDB test sets. (a)∼(b) The violin plots of precisions of the top 50 predicted contacts by each method on (a) HomoPDB and (b) HeteroPDB. The mean precisions of the predictions are highlighted in green lines, and the median precisions of the predictions are highlighted in gray lines. (c)∼(d) The complementary cumulative distribution function of precisions of the top 50 predicted contacts by each method on (c) HomoPDB and (d) HeteroPDB.

We further analyzed the performance of DRN-1D2D_Inter on PPI targets with various sizes and contact densities. Specifically, we grouped targets in the two test sets according to the mean length 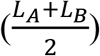 the contact density 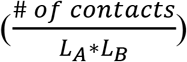 of the two interacting proteins of each PPI. In **Figure 3**, we show mean precisions of the top 50 contacts predicted by different models on PPIs within different intervals of mean lengths and contact densities. As we can see from Figure 3, DRN-1D2D_Inter still consistently outperformed the other two models on targets with different sizes and contact densities, which illustrate the robustness of DRN-1D2D_Inter. Besides, we can also see from the figure that all the methods tend to achieve better performances when the target PPI has higher contact densities, but the performance does not have a clear dependency on the PPI size. It is reasonable since it is obviously more challenging to identify the contacting ones when the residue pairs have lower contact probabilities.

**Figure 3.**
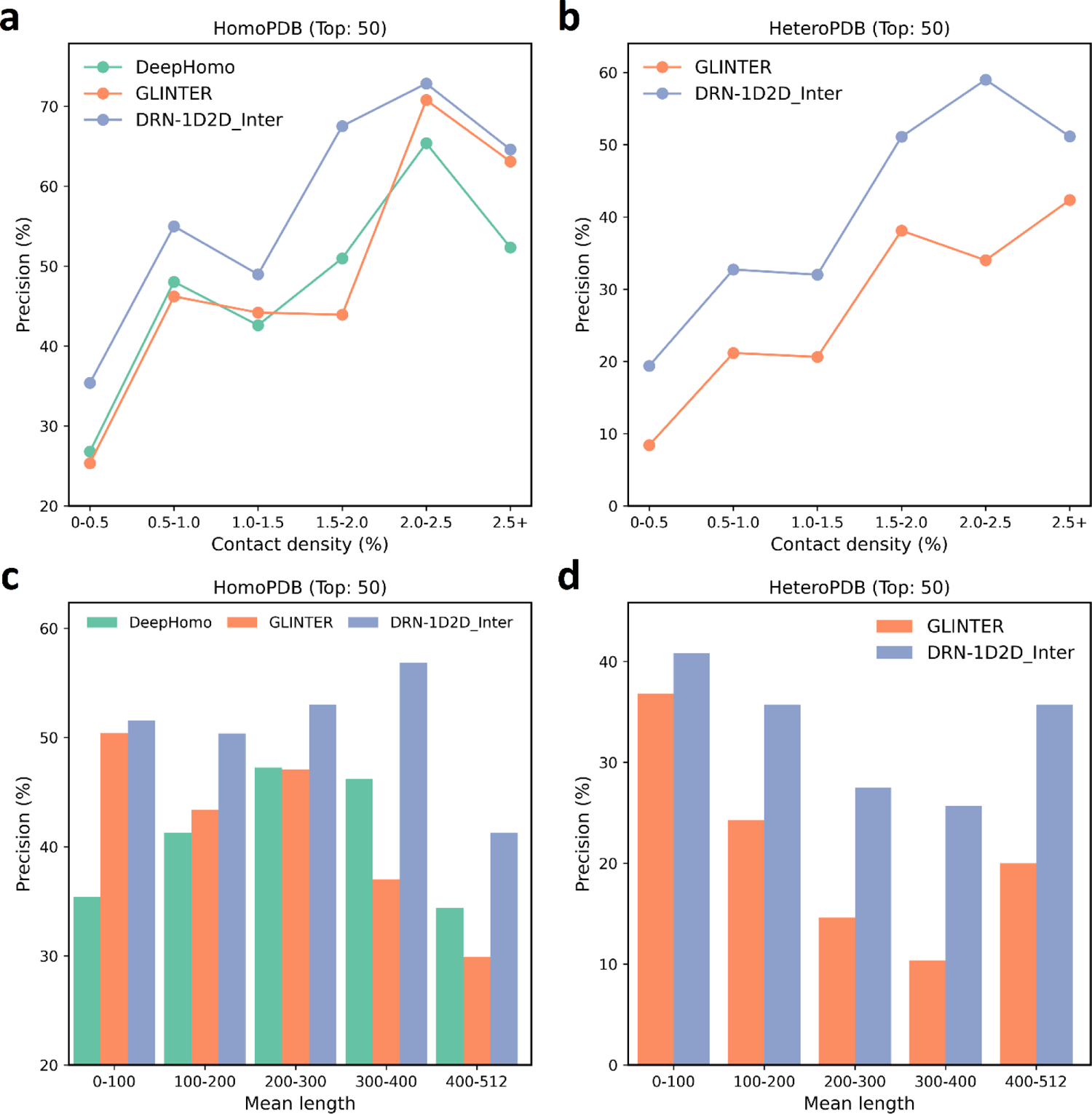
Performances of inter-protein contact prediction by different methods on PPIs with various sizes and contact densities in the HomoPDB and HeteroPDB test sets. (a)∼(b) The mean precisions of the top 50 contacts predicted by each method on PPIs within different intervals of contact densities in (a) HomoPDB and (b) HeteroPDB. (c)∼(d) The mean precisions of the top 50 contacts predicted by each method on PPIs within different intervals of mean lengths in (c) HomoPDB and (d) HeteroPDB.

### 3. The performance of DRN-1D2D_Inter on the DHTest and DB5.5

We further evaluated all the six methods on DHTest and BM5.5. Still DeepHomo was only evaluated on DHTest, as it can only be used to predict inter-protein contacts for homomeric PPIs. In Table 2, we show the mean precisions of the top (5, 10, 50, L/10, L/5) predicted contacts (L as the sequence length of the shorter protein in the PPI) by the six methods on the two test sets (Note: GLINTER failed to produce valid prediction results for 47 targets in DHTest and 3 targets in DB5.5, thus we removed these targets in the evaluation of the performance of GLINTER. The performances of other methods on HomoPDB and HeteroPDB after the removal of these targets are shown in Table S2). As we can see from Table 2 that DNR-1D2D_Inter still consistently achieved the best performance among these models. It can also be found from the table that the performance of DRN-1D2D_Inter on the hetero PPI dataset DB5.5 is significantly lower than that on HeteroPDB, and so are the performances of all other five methods. One reason might be the mean contact density of PPIs in DB5.5 is significantly lower than that in HeteroPDB (see Table S1). We further performed more detailed analysis on the performance comparison between DRN-1D2D_Inter and the two state-of-the-art methods GLINTER and DeepHomo. Specifically, the violin plots and the complementary cumulative distribution function of precisions of the top 50 predicted contacts by each method are shown in Figure 4a-b and Figure 4c-d respectively. It can be clearly seen from the figure that that DRN-1D2D_Inter outperformed DeepHomo (p-value: 6.11e-2) and GLINTER (DHTest p-value: 9.51e-9, DB5.5 p-value: 1.95e-2).

**Table 2.** The performances of CCMpred, ESM-1b, ESM-MSA-1b, DeepHomo, GLINTER and DRN-1D2D_Inter on the DHTest and DB5.5 test sets.

| Methods | DHTest |  |  |  |  | DB5.5 |  |  |  |  |
| --- | --- | --- | --- | --- | --- | --- | --- | --- | --- | --- |
|  | L/5 | L/10 | 50 | 10 | 5 | L/5 | L/10 | 50 | 10 | 5 |
| CCMpred | 3.22 | 3.81 | 3.28 | 3.77 | 4.15 | 1.52 | 2.19 | 1.73 | 2.20 | 2.03 |
| ESM-1b | 6.87 | 7.18 | 6.65 | 6.78 | 6.62 | 1.19 | 1.11 | 1.02 | 1.19 | 1.36 |
| ESM-MSA-1b | 8.08 | 7.98 | 7.62 | 8.54 | 8.77 | 0.77 | 0.56 | 0.78 | 0.85 | 0.34 |
| DeepHomo | 45.0 | 47.4 | 44.2 | 50.2 | 50.9 |  |  |  |  |  |
| GLINTER | 48.5 | 50.8 | 47.4 | 51.2 | 52.3 | 18.8 | 20.5 | 15.2 | 21.9 | 20.7 |
| DRN-1D2D_Inter | **52.1** | **53.3** | **52.3** | **54.9** | **56.0** | **24.4** | **26.9** | **20.7** | **25.4** | **27.5** |

Note: 1. The highest mean precision in each column is highlighted in bold.

**Figure 4.**
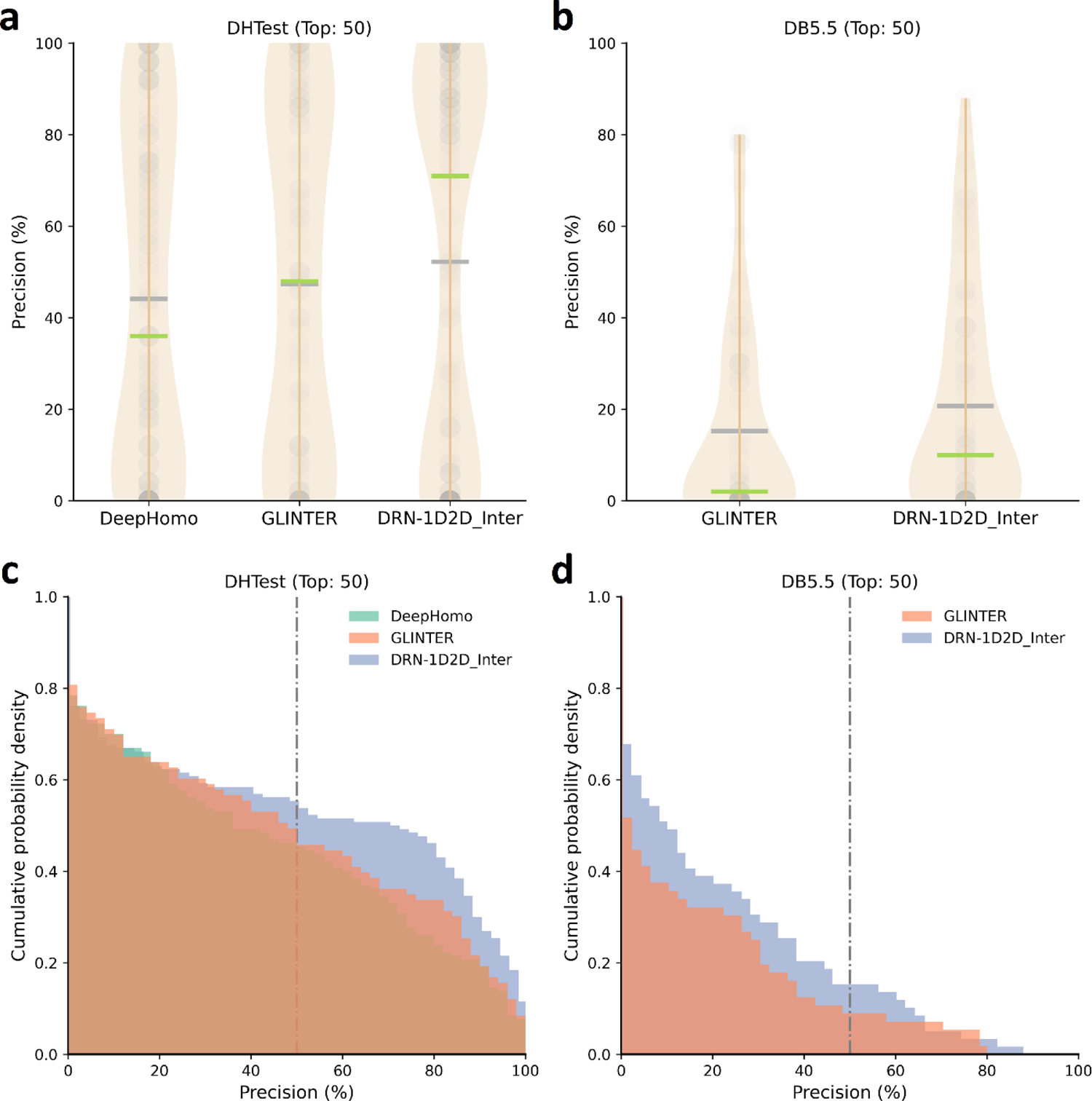
The statistical analysis of precisions of the top 50 predicted contacts by different methods on the DHTest and DB5.5 test sets. (a)∼(b) The violin plots of precisions of the top 50 predicted contacts by each method on (a) DHTest and (b) DB5.5. The mean precisions of the predictions are highlighted in green lines, and the median precisions of the predictions are highlighted in gray lines. (c)∼(d) The complementary cumulative distribution function of precisions of the top 50 predicted contacts by each method on (c) DHTest and (d) DB5.5.

### 4. Ablation study

To evaluate the contributions of different input components to the model performance, we conducted ablation study on the model of DRN-1D2D_Inter. As the input features of DRN-1D2D_Inter can be grouped into four categories: (1) the 1D features from sequences of the interacting protein (ESM-1b embedding), (2) the 1D features from the MSAs of homologous sequences of the interacting protein (PSSM + ESM-MSA-1b embedding), (3) the 2D features from the paired sequence (ESM-1b attention), (4) the 2D features from the paired MSA (CCMpred + alnstats + ESM-MSA-1b attention), we trained four additional models with the application of different combinations of the four categories of the input features of DRN-1D2D_Inter as their input features. Specifically, with the application of feature category 1) as the baseline, we successively included the three additional feature categories in the models (model a: feature (1); model b: feature (1) + (2); model c: feature (1) + (2) +(3); model d: feature (1) + (2) + (3) +(4)). All the four models were trained using the same protocol as DRN-1D2D_Inter on the same training and validation partition without cross validation. We further evaluated performances of four models together with DRN-1D2D_inter (model e: model d + cross validation) on HomoPDB and HeteroPDB respective. In Table 3, we show the mean precisions of the top (5, 10, 50, L/10, L/5) predicted contacts (L as the sequence length of the shorter protein in the PPI) by the five models on the two test sets. Besides, the performances of DeepHomo and GLINTER are also included in the table for reference. As it can be seen from the table that our baseline model has already yielded a performance not far from the performances of DeepHomo and GLINTER, although our baseline model only used the sequence embeddings as the input features. It can also be seen from the table both the inclusion of additional input features and the cross validation improved the model performance. In Figure 5a, we show the increments of the mean precisions of the top 50 predicted contacts by different ablation study models over the baseline model (model a) on the two datasets respectively. We can clear see from the figure that apart from the sequence embeddings, the 2D features from the paired MSA also play very important role in the model performance. In Figure 5b-c, we show the performance comparison (top 50 predictions) between model c and model d for each target in HomoPDB and HeteroPDB respectively (see Figure S1 for model a (baseline) versus model e (DRN-1D2D_Inter)). It can be clearly seen from Figure 5b-c that the inclusion of 2D features from the paired MSA significantly improved the model performance (HomoPDB p-value: 3.04e-6, HeteroPDB p-value: 1.02e-2).

**Table 3.**
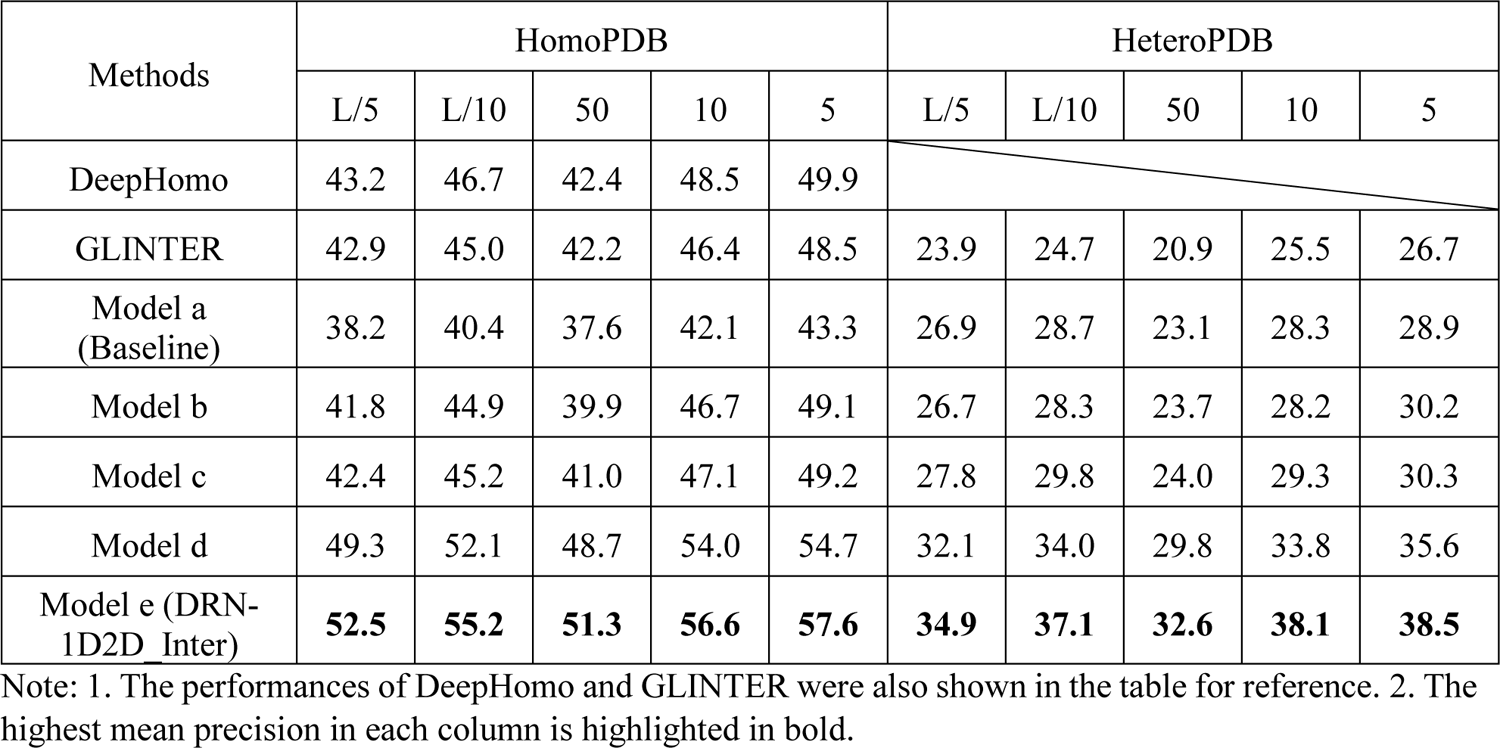
The performances of different ablation study models on the HomoPDB and HeteroPDB test sets.

**Figure 5.**
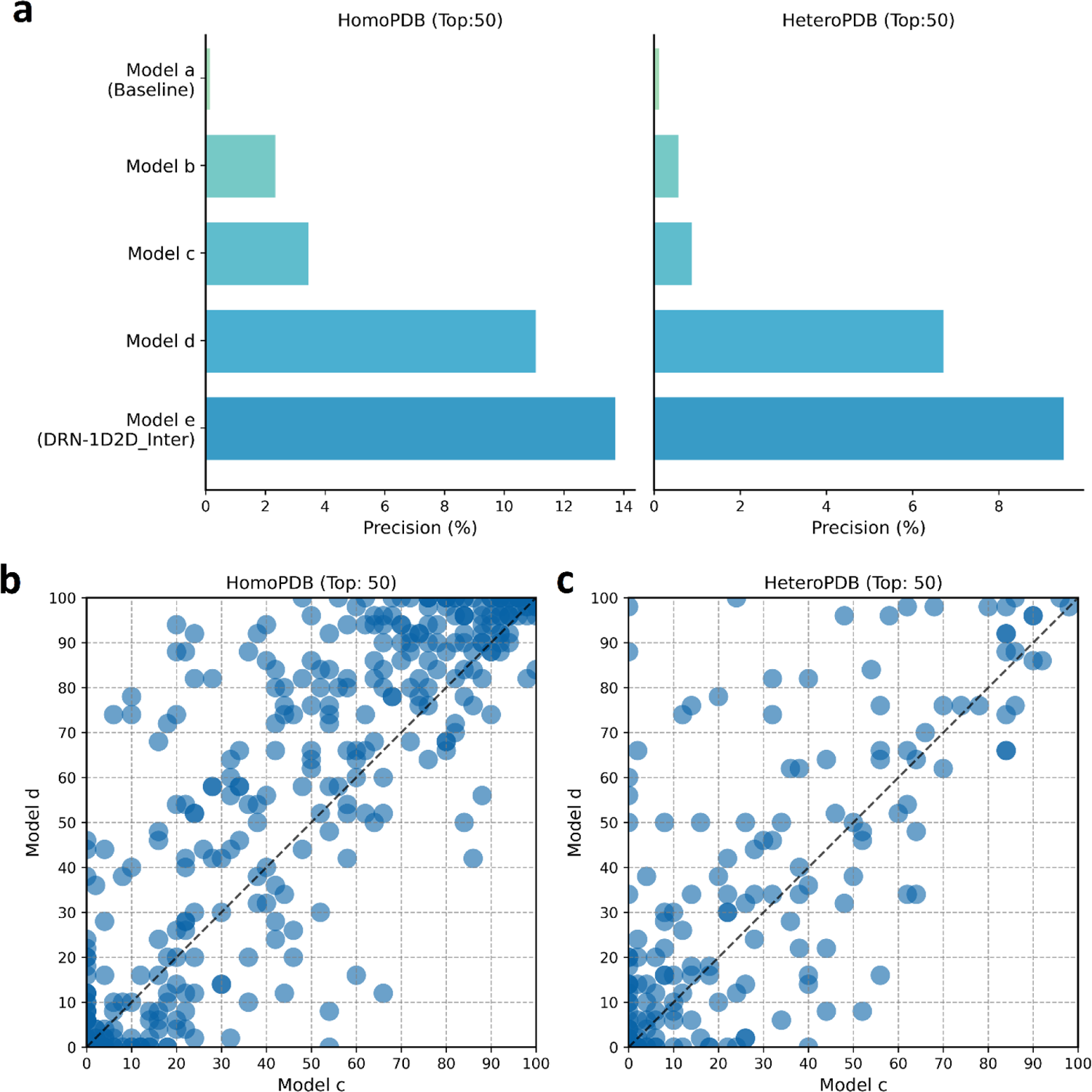
Ablation study of DRN-1D2D_Inter on the HomoPDB and HeteroPDB test sets. (a) The increments of the mean precision of the top 50 predicted contacts by different ablation study models over the baseline model (model a) on HomoPDB and HeteroPDB. (b)∼(c) The comparison of the precisions of the top 50 predicted inter-protein contacts by model c and model d for each target in (b) HomoPDB and (c) HeteroPDB.

### 5. Single sequence-based inter-protein contact prediction

As it is shown in the ablation study that the baseline model can achieve a performance not far from the performances of the two state-of-the-art methods DeepHomo and GLINTER. The baseline model only requires single sequence embeddings as the input, which is extremely computationally efficient, since it does not require the building and pairing of MSAs. Accurate inter-protein contact prediction from single sequence is attractive, since it would easily open the door to genome-wide PPI prediction and inter-protein contact prediction. To further improve the performance of the single sequence inter-protein contact prediction, we trained seven independent single sequence models through cross validation, and the final model is the average of the seven models, which is referred to as SS_Inter. We further evaluated SS_Inter together with DeepHomo and GLINTER on the four test sets used in our study. In Table 4, we show the mean precisions of the top (50, L/5) predicted contacts (L as the sequence length of the shorter protein in the PPI) by DeepHomo, GLINTER and SS_Inter on the four test sets: HomoPDB, HeteroPDB, DHTest and DB5.5. From the table we can clearly see that except for DHTest, SS_Inter outperformed GLINTER and DeepHomo on all other test sets (see Figure S2 for the performance comparison for each target in the four datasets). It is encouraging that the single sequence model can achieve comparable performance to the two state-of-the-art models, considering the latter models employed very complex input features.

**Table 4.**
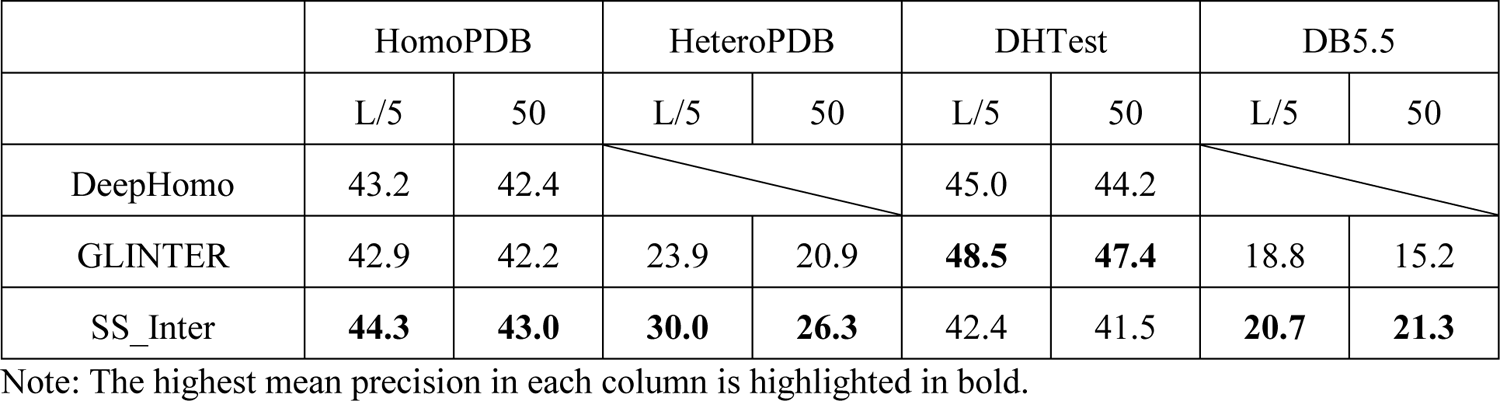
The performances of DeepHomo, GLINTER and SS_Inter on the HomoPDB, HeteroPDB, DHTest and DB5.5 test sets.

## Discussion

Predicting contacting residue pairs between interacting proteins is of great significance, as the knowledge of inter-protein residue-residue contacts is very useful for structural characterization of PPIs. However, accurate inter-protein contact prediction requires the identification of tens of contacting residue pairs from hundreds of thousands of candidates, which is extremely challenging, and performances of existing inter-protein contact prediction methods are quite limited. In this study, we developed a deep learning method for inter-protein contact prediction, referred to as DRN-1D2D_Inter. DRN-1D2D_Inter mainly employed structural information enriched input features produced by two protein language models (ESM-1b and ESM-MSA-1b), which are then transformed by dimensional hybrid residual networks to predict inter-protein contacts. Comparing with previous methods, DRN-1D2D_Inter has two major innovations: First, we employ protein language models to generate structural information enriched features rather than extracting structural features directly from monomeric structures, which allows us to make inter-protein contact prediction purely from sequences; second, we leverage dimensional hybrid residual blocks rather than traditional 2D residual blocks to build the residual networks, and our previous study shows that application of the dimensional hybrid residual blocks can increase the effective receptive field of the network, and thus improve the model performance. Extensively benchmarked DRN-1D2D_Inter on multiple datasets including both homomeric and heteromeric PPIs, we show DRN-1D2D_Inter consistently and significantly outperforms GLINTER and DeepHomo, the two state-of-the-art inter-protein contact prediction methods. It is worth noting that DRN-1D2D_Inter made the prediction purely from sequences, but the latter methods leveraged native structures of the interacting proteins in the prediction. Further ablation study shows that the single sequence embeddings play a critical role in the model performance. Due to its extremely high computational efficiency, we further developed single sequence-based inter-protein contact prediction model referred to as SS_Inter. Extensive benchmarking result shows that SS_Inter can achieve comparable performance to GLINTER and DeepHomo. The development of accurate single sequence-based inter-protein contact prediction methods would open the door to genome-wide PPI prediction and inter-protein contact prediction.

## Availability

The code and data for DRN-1D2D_Inter can be found through https://github.com/ChengfeiYan/DRN-1D2D_Inter.

## Acknowledgement

The work was supported by the National Natural Science Foundation of China (32101001) and new faculty startup grant (3004012167) of Huazhong University of Science and Technology.

## Methods

### 1. The training and test datasets preparation

#### 1.1 The preparation of the non-redundant PPI dataset

##### 1.1.1 Collecting non-redundant homomeric PPIs

To collect non-redundant homomeric PPIs, we first filtered the structures deposited in Protein Data Bank (PDB)^6^ before March 7, 2022 according to the following criteria: 1) Experimental Method equals to X-RAY DIFFRACTION; 2) Data Collection Resolution ≤ 3Å; 3) Entry Polymer Composition equals to homomeric protein; 4) Total Number of Polymer Instance (Chains) equals to 2; 5) Symmetry Symbol equals to C2; 6) Entry Polymer Types equals to Protein (only). A total of 19446 PDB entries were obtained. We then removed entries that have more than one assembly, ensuring each PPI has a unique binding mode. The remaining entries were further filtered with the following criteria: 1) The length of each chain is not shorter than 30 and not longer than 512. 2) the number of inter-protein contacts is larger than 10. 16349 entries of homomeric PPIs were kept at this step. It should be noted that an inter-protein contact is defined if the 3D distance of any heavy atoms of two residues from the two interacting proteins is less than or equal to 8 Å. Here the chain length of the protein monomer was restricted to be not longer than 512, for the longest sequence length allowed by ESM-MSA-1b is 1024.

To remove redundancy, we clustered the 16349 sequences from the 16349 homomeric PPIs using CD-HIT^25^ with 40% as the sequence similarity threshold. In each cluster, we only kept the PPI with the largest number of inter-protein contacts. Finally, 4928 non-redundant homomeric PPIs were obtained.

##### 1.1.2 Collecting the non-redundant heteromeric PPIs

To collect non-redundant heteromeric PPIs, we first filtered the structures deposited in Protein Data Bank (PDB) before March 7, 2022 according to the following criteria: 1) Experimental Method equals to X-RAY DIFFRACTION; 2) Data Collection Resolution ≤ 3A; 3) Entry Polymer Composition equals to heteromeric protein; 4) Total Number of Polymer Instance (Chains) greater than or equal to 2; A total of 23603 entries were obtained. We then removed entries that have more than one assembly, ensuring that each PPI has a unique binding mode. Then protein chains in each entry were paired to form protein pairs, and in total of 23551 protein pairs were obtained. The same as the case of homomeric PPIs, we further filtered the protein pairs with the following citeria: 1) The length of each chain is not shorter than 30 and not longer than 512. 2) the number of inter-protein contacts is larger than 10. 13275 heteromeric PPIs were kept at this step.

To remove redundancy, we clustered 26550 sequences from 13275 heteromeric PPIs using CD-HIT with 40% as sequence similarity threshold. Each sequence is then uniquely labeled with the cluster (e.g. cluster 0, cluster 1, …) to which it belongs, from which each heteromeric PPI can be marked with a pair of cluster IDs (e.g. cluster 0-cluster 1). The PPIs belonging to the same cluster pair (note: cluster n - cluster m and cluster n-cluster m were considered as the same pair) were considered as redundant, for which we only kept the PPI with largest number of inter-protein contacts. 3134 non-redundant heteromeric PPIs were obtained.

#### 1.2 The preparation of DHTest

The DH test set was formed by removing PPIs redundant to our training dataset from the original test set of DeepHomo, which contains 300 homomeric PPIs. Specifically, the 300 monomer sequences from the 300 homomeric PPIs were first merged with the 10296 (4428+2934*2) sequences from our training set, and then all the 13286 sequences were clustered using CD-HIT with 40% as the sequence similarity threshold. Each sequence was uniquely labeled by the cluster to which it belongs, and then each PPI was labeled with a pair of clusters (cluster n-cluster m, for homomeric PPIs, n always equals to m). For the 300 PPIs in the test set of DeepHomo, we only kept PPIs which did not share the cluster pair with any PPI in our training set. Finally, 130 homomeric PPIs were kept in the DHTest test set.

#### 1.3 The preparation of DB5.5

The BM5.5 test set was formed by removing PPIs redundant to our training dataset from the heterodimers in Protein-protein Docking Benchmark 5.5. The Protein-protein Docking Benchmark contains a total of 257 targets, of which 153 are heterodimers. Specifically, the 306 sequences from the 153 heterodimers were first merged with the 10296 (4428+2934*2) sequences from our training set, and then all the 10602 sequences were clustered using CD-HIT with 40% as the sequence similarity threshold. Each sequence is uniquely labeled by the cluster to which it belongs, and then each PPI was labeled with a pair of clusters (note: cluster n-cluster m and cluster n-cluster m were considered as the same pair). For the 153 heteromeric PPIs, we only kept PPIs which did not share the cluster pair with any PPI in our training set. Finally, 59 heteromeric PPIs were kept in the DB5.5 test set.

### 2. The preparation of input features

The 1D sequence embeddings of the two interacting proteins were directly obtained by inputting each protein sequence to ESM-1b separately. To obtain the 2D inter-protein attention maps from the paired sequence, we inputted the paired sequence to ESM-1b to generate the attention maps including both the intra- and inter-protein pairwise attentions, from which we extracted the inter-protein attention maps.

To obtain input features drawn from the MSA for each interacting protein and the paired MSA, the MSAs and the paired MSA should be prepared first. Specifically, Jackhammer was applied to search the UniRef100/UniRef90 database^26^ (UniRef100 was used to build MSAs for the test set and Uniref90 was used for the training set to save the computational time) to build MSA for each sequence. For homomeric PPIs, we only prepared MSA for one of the two interacting proteins, as sequences of the two interacting proteins are the same. The paired MSA for each homomeric PPI was formed by concatenating two copies of the MSA. For heteromeric PPIs, the paired MSA was formed by pairing the MSAs through the phylogeny-based approach described in ^27^.

To obtain the PSSM for each interacting protein, the corresponding MSA was inputted to hhmake^28^ to get the HMM file, and then the LoadHMM.py script from RaptorX_Contact^29^ was used to extract the PSSM. Since ESM-MSA-1b requires huge memory of GPU when the input MSA contains two many sequences, in this study, to obtain the MSA embedding for each interacting protein, the number of sequences in the MSA was limited to 256 through the MSA downsampling with hhfilter using parameter “-diff 256”, which maximized the diversity of the kept sequences, as it was suggested in the literature of ESM-MSA-1b. The downsampled MSA was then used as the input to ESM-MSA-1b to generate the MSA embedding.

The evolutionary coupling matrix was obtained by inputting the paired MSA to CCMpred with its default settings. The mutual information matrix, the APC-corrected mutual information matrix and contact potential matrix were obtained by inputting the paired MSA to alnstats. To obtain the attention maps using ESM-MSA-1b, the paired MSA was also downsampled with hhfilter^28^ using parameter “-diff 256” to limit the paired sequence in the MSA to 256, which was then inputted to ESM-MSA-1b to generate the attention maps. All the 2D pairwise features drawn from the paired MSA included intra- and inter-protein pairwise features. In this study, only the inter-protein portions were extracted to form the 2D input features of our model.

### 3. The implementation of reference methods and statistical tests

CCMpred (https://github.com/soedinglab/CCMpred) was implemented with its default settings. ESM-1b and ESM-MSA-1b was implemented through the provide Evolutionary Scale Modeling (ESM) package (https://github.com/facebookresearch/esm). DeepHomo was implemented directly through the provided webserver (http://huanglab.phys.hust.edu.cn/deephomo/). GLINTER was implemented through the provided package (https://github.com/zw2x/glinter). It should be noted that GLINTER sometimes failed to provide prediction results for unknown reasons (most of the failed cases are homomeric PPIs). The p values for performance comparison were calculated through Mann-Whitney U rank test.

