## Supplemental PDF for "Improved inter-protein contact prediction using dimensional hybrid residual networks and protein language models"

### Supplementary

#### Supplemental Tables and Figures

**Table S1. The statistics of training set, HomoPDB, HeteroPDB, DHTest and DB5.5**

| | Training set | HomoPDB | HeteroPDB | DHTest | DB5.5 |
|---|---|---|---|---|---|
| Target number | 7362 | 400 | 200 | 130 | 59 |
| Mean contact density (%) | 1.28 | 1.33 | 1.29 | 1.448 | 1.01 |
| Median contact density (%) | 0.698 | 0.783 | 0.753 | 0.923 | 0.67 |
| Mean min length | 208 | 239 | 150 | 250 | 136 |
| Median min length | 187 | 220 | 135 | 219 | 123 |
| Mean paired length | 470 | 479 | 426 | 501 | 424 |
| Median paired length | 436 | 440 | 391 | 438 | 401 |

**Table S2. The performances of DeepHomo, GLINTER and DRN-1D2D_Inter on HomoPDB and HeteroPDB after the removal of targets which GLINTER failed to make the prediction**

| Methods | HomoPDB | | | | | HeteroPDB | | | | |
|---|---|---|---|---|---|---|---|---|---|---|
| | L/5 | L/10 | 50 | 10 | 5 | L/5 | L/10 | 50 | 10 | 5 |
| CCMpred | 3.45 | 3.17 | 4.17 | 3.14 | ·3.07 | 5.96 | 7.00 | 4.65 | 7.14 | 8.54 |
| ESM-1b | 6.15 | 6.44 | 5.52 | 6.55 | 6.83 | 2.56 | 2.61 | 3.30 | 2.49 | 2.60 |
| ESM-MSA-1b | 5.62 | 5.84 | 5.47 | 5.64 | 5.46 | 7.61 | 7.84 | 6.62 | 8.90 | 8.22 |
| DeepHomo | 41.1 | 44.3 | 40.6 | 46.0 | 46.9 | | | | | |
| GLINTER | 42.8 | 45.0 | 42.2 | 46.4 | 48.5 | 23.9 | 24.7 | 20.9 | 25.5 | 26.7 |
| DRN-1D2D_Inter | **50.3** | **53.0** | **49.3** | **54.0** | **54.9** | **33.9** | **35.9** | **31.6** | **36.5** | **37.2** |

Note: The highest mean precision in each column is highlighted in bold.

**Table S3. The performances of DeepHomo, GLINTER and DRN-1D2D_Inter on DHTest and DB5.5 after the removal of targets which GLINTER failed to make the prediction**

| Methods | DHTest | | | | | DB5.5 | | | | |
|---|---|---|---|---|---|---|---|---|---|---|
| | L/5 | L/10 | 50 | 10 | 5 | L/5 | L/10 | 50 | 10 | 5 |
| CCMpred | 3.96 | 4.69 | 3.47 | 5.06 | 5.31 | 1.40 | 2.11 | 1.68 | 1.96 | 2.14 |
| ESM-1b | 5.85 | 6.62 | 5.76 | 6.02 | 6.51 | 1.25 | 1.17 | 1.07 | 1.25 | 1.43 |
| ESM-MSA-1b | 6.99 | 7.36 | 6.99 | 7.47 | 8.43 | 0.81 | 0.59 | 0.82 | 0.89 | 0.36 |
| DeepHomo | 41.6 | 43.9 | 41.3 | 47.6 | 48.7 | | | | | |
| GLINTER | **48.5** | **50.7** | **47.4** | **51.2** | **52.2** | 18.8 | 20.5 | 15.2 | 21.9 | 20.7 |
| DRN-1D2D_Inter | 45.8 | 47.2 | 45.9 | 49.5 | 50.8 | **23.8** | **26.0** | **19.6** | **24.6** | **27.1** |

Note: The highest mean precision in each column is highlighted in bold.

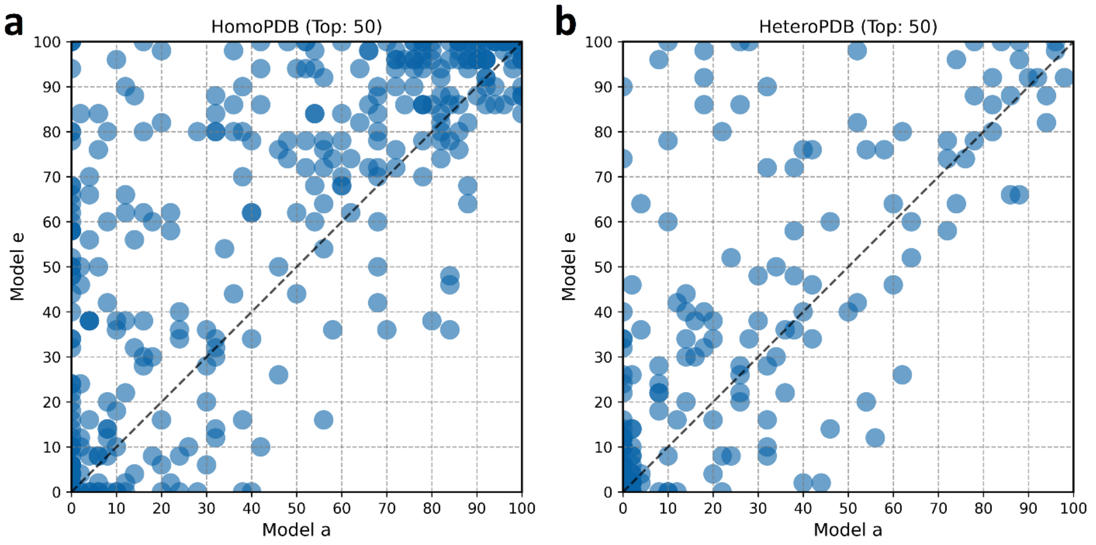


**Fig S1.** The comparison of the precisions of the top 50 predicted inter-protein contacts by model a (the baseline model) and model e (DRN-1D2D_Inter) for each target in (b) HomoPDB and (c) HeteroPDB.

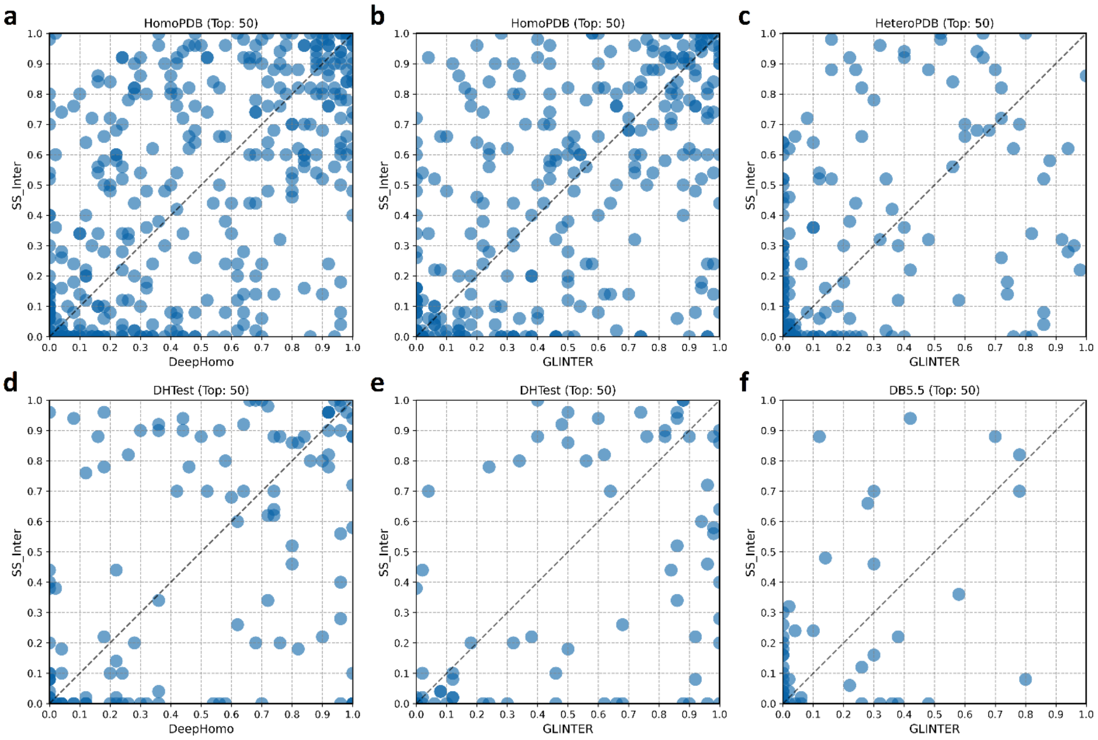


**Fig S2.** The comparison of the precisions of the top 50 predicted inter-protein contacts by SS_Inter, DeepHomo and GLINTER for each target in HomoPDB, HeteroPDB, DHTest and DB5.5. (a) SS_Inter versus DeepHomo on HomoPDB. (b) SS_Inter versus GLINTER on HomoPDB. (c) SS_Inter versus GLINTER on HeteroPDB. (d) SS_Inter versus DeepHomo on DHTest. (e) SS_Inter versus GLINTER on DHTest. (f) SS_Inter versus GLINTER on DB5.5.
